# Cell-APP: A generalizable method for cell annotation and cell-segmentation model training

**DOI:** 10.1101/2025.01.23.634498

**Authors:** Anish J. Virdi, Ajit P. Joglekar

**Affiliations:** Department of Biophysics, University of Michigan, Ann Arbor, MI 48109, USA; Cell & Developmental Biology, University of Michigan Medical School, Ann Arbor, MI 48109, USA

## Abstract

Deep learning-based segmentation models can accelerate the analysis of high-throughput microscopy data by automatically identifying and classifying cells in images. However, the datasets needed to train these models are typically assembled via laborious hand-annotation. This limits their scale and diversity, which in turn limits model performance. We present Cell-APP (Cellular Annotation and Perception Pipeline), a tool that automates the annotation of high-quality training data for transmitted-light (TL) cell segmentation. Cell-APP uses two inputs—paired TL and nuclear fluorescence images—and operates in two main steps. First, it extracts each cell’s location from the nuclear fluorescence channel and provides these locations to promptable deep learning models to generate cell masks. Then, it classifies each cell as mitotic or non-mitotic based on nuclear features. Together, these masks and classifications form the basis for cell segmentation training data. By training vision-transformer-based models on Cell-APP-generated datasets, we demonstrate how Cell-APP enables the creation of both cell line-specific and multi-cell line segmentation models. Cell-APP thus empowers laboratories to tailor cell segmentation models to their needs, and outlines a scalable path to creating general models for the research community.

**Significance Statement:**

1. Deep learning-based cell segmentation models are typically trained on manually annotated datasets. Manual annotation limits dataset scalability and, consequently, the ability of trained models to generalize across cell types.
2. Cell-APP automates mask generation and cell classification to rapidly create large, custom training datasets. It uses Meta AI’s SAM for mask generation and extracts information from fluorescence signals for classification.
3. By reducing the need for hand-annotation, Cell-APP lowers the cost associated with building cell line-specific and generalist segmentation models. It may accelerate high-throughput image analysis and democratize model development across research labs.

## Introduction

High-throughput, time-lapse, live-cell microscopy enables researchers to collect large, spatially and temporally resolved datasets that capture the behaviors of thousands of cells. These datasets, once analyzed, provide the statistical power needed to model and understand complex, heterogeneous cellular phenomena. To analyze these data, one must identify, segment, and in some cases classify every cell in the dataset. If performed manually, these tasks, collectively known as instance segmentation, are time-consuming and severely limit data output. These motivating factors have led to the development of many algorithms that perform instance segmentation of cells in 2-D tissue cultures. Of these algorithms, deep learning models define the current state-of-the-art (Falk *et al*., 2019; Patel *et al*., 2019; Fazeli *et al*., 2020; Cohen and Uhlmann, 2021; Fishman *et al*., 2021). These models can be sorted into two categories by use case: those that perform best on fluorescent images, such as Aura-Net, and those that perform best on transmitted-light (TL) images, such as the ones produced by LIVECell and EVICAN (Carpenter *et al*., 2006; Schmidt *et al*., 2018; Rivenson *et al*., 2019; Ling *et al*., 2020; Edlund *et al*., 2021; Stringer and Pachitariu, 2025). CellPose and StarDist straddle both categories. These algorithms have transformed how researchers analyze microscopy data.

Despite the success of fluorescence segmentation models, in cell biology studies, the fluorescence signal of interest may not be suitable for segmentation. For example, the protein Cyclin B is all but completely degraded at the end of mitosis (Clute and Pines, 1999). As a result, segmentation based on Cyclin B fluorescence would fail to capture cells in telophase and early G1, preventing lineage tracing. To overcome this, researchers may use an additional constitutively expressed fluorophore for cell segmentation. However, this approach consumes a channel that could otherwise provide biological information. Moreover, fluorophores used for segmentation may undergo photobleaching; this will cause segmentation quality to change throughout time-lapse experiments.

These problems can be avoided by segmenting TL images instead. This task, however, is not without challenges. Chief among them is that morphology varies drastically across cell lines and imaging setups; this makes it difficult to create a comprehensive training dataset. Such a dataset is desirable, as it could be used to train a model that performs well across diverse morphologies—a capability that current models lack (Stringer *et al*., 2021). Researchers have employed teams of hand annotators in efforts to create such a morphology-diverse dataset. The LIVECell and EVICAN datasets, which respectively contain over 1.6 million and 52,000 annotated cells, were created in this way (Schwendy *et al*., 2020; Edlund *et al*., 2021). Still, encapsulating the full array of morphologies displayed by the over 1,500 animal cell lines offered by American Type Culture Collections is an open challenge. It may also be desirable to create training datasets specific to the imaging modality (wide-field, differential interference contrast, etc.) and magnification. Finally, these datasets cannot be used to train a model that simultaneously classifies cells (e.g., mitotic vs. non-mitotic), an ability of great use for biologists. For these reasons, we must augment these datasets or create new ones to develop improved deep learning-based cell segmentation models.

Here, we present Cell-APP (Cellular Annotation and Perception Pipeline), a tool that automates cellular mask generation and cell classification to create custom instance segmentation training datasets in TL microscopy. With Cell-APP-generated datasets, users can train segmentation models tailored to their cell lines and imaging conditions. The tool requires only two inputs—paired TL and nuclear fluorescence images—and reliably segments and classifies the majority of cells present. Using Cell-APP, we created expansive datasets of HeLa, U2OS, HT1080, and RPE-1 cells and used them to train both cell line-specific and general-purpose vision-transformer-based models for TL cell segmentation.

## Results

### Automating mask generation with µSAM

A tool that automates cellular mask generation must be able to: (1) find each cell in an image and (2) generate corresponding polygonal masks. One challenge regarding (2) is that cells are incredibly diverse in appearance, and their appearance depends on the setup with which they are imaged. Therefore, for a tool to succeed at (2), it must be capable of generating masks for a wide variety of objects in various contexts—including types of objects it has not seen before. In computer vision, this capability is called zero-shot generalization.

Meta AI’s Segment Anything Model (SAM) meets most of these requirements. SAM is a deep learning-based instance segmentation model inspired by large language models (LLMs) (Kirillov *et al*., 2023; Archit *et al*., 2025). LLMs have demonstrated strong zero-shot generalization (Brown *et al*., 2020). Much of their success stems from prompt engineering, in which human-generated text influences the LLM to generate a valid response for the task at hand. Meta AI designed SAM to similarly learn from prompts, but in visual contexts. It encodes user-provided centroids, bounding boxes, masks, or text alongside the input image during training and inference. These prompts enhance its understanding of “objects” and enable it to generate accurate masks for them. We used a biologically fine-tuned version of SAM, called *μ*SAM (Archit *et al*., 2025). µSAM excels at generating cell masks if user-generated prompts are provided. Thus, µSAM satisfies (2) but not (1).

To fully automate mask generation, we programmatically generated markers for all cells in each TL image and used them to prompt µSAM. Monolayered tissue culture cells have nuclei that are usually centrally located and non-overlapping. Cell-APP uses these nuclei to prompt µSAM in the following manner: given a TL and nuclear fluorescence image pair, it computes the centroid and bounding box of each nucleus using digital image processing techniques (**Methods**). It then provides either the centroids or bounding boxes to µSAM alongside the TL image during inference (**Figure 1**). This process automates mask generation.

**Figure 1.**
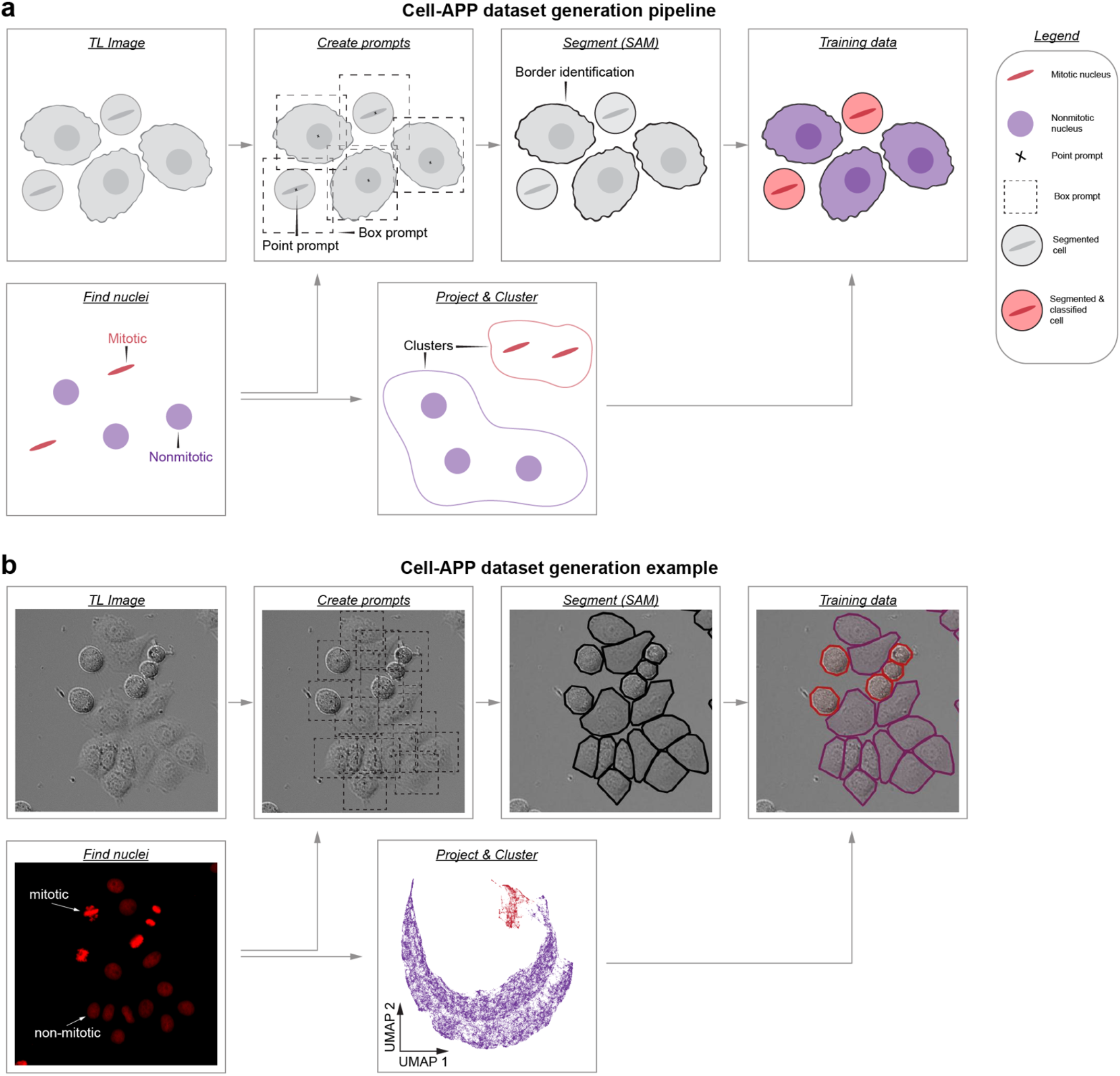
Cell-APP automates the generation of cell-segmentation datasets. **a**, Cartoon schematic of the dataset generation pipeline. The pipeline uses digital image processing techniques to find nuclei, compute their centroids, and determine a bounding box size *(bottom left)*. It then overlays nuclei and bounding boxes on the TL image *(top; second from left)* and feeds the TL image/prompt combination to a Segment Anything model for segmentation *(top; third from left)*. In parallel, computed bounding boxes are used to crop single nuclei from the chromatin image. Geometric and intensity-based features are computed for each nucleus and used to classify them as mitotic or non-mitotic *(bottom right)*. Classifications are then merged with segmentations to form the dataset *(top right)*. **b**, Microscopy-based schematic of **a**.

### Automating classification with chromatin morphology

Having automated mask generation, we sought a method to automatically classify each mask as either non-mitotic or mitotic. Previous work has demonstrated that neural networks can be trained to classify images of single nuclei as either non-mitotic or mitotic (Ulicna *et al*., 2021). This is possible as chromatin undergoes drastic topological changes when cells enter mitosis. In non-mitotic metazoan cells, chromatin appears as an ovoid of uniform intensity. As the cell transitions to mitosis, the chromatin condenses into scattered chromosomes (prophase). These chromosomes then align into a thin, intense line (metaphase), before the mitotic spindle segregates them into two daughter cells (anaphase). All these phases are visually distinct (**Figure 1b**).

We therefore developed an automated procedure to classify nuclei and map those classifications to the corresponding SAM-generated masks. The procedure first uses the computed bounding boxes to crop nuclei from the nuclear fluorescence images. Then, for each nucleus, it computes geometric and intensity-based properties; these properties form a high-dimensional vector that characterizes the nucleus (see **Table 1** for a list of properties). It then uses the Uniform Manifold Approximation and Projection (UMAP) algorithm to project these vectors onto the plane, where they can be clustered (McInnes *et al*., 2018). For clustering, it leverages the Hierarchical Density-based Spatial Clustering of Applications with Noise (HDBSCAN) algorithm (**Figure 1b**) (Campello *et al*., 2013). With this approach, we can accurately classify nuclei as mitotic or non-mitotic.

### Cell line-specific datasets

These solutions to mask generation and cell classification, when applied sequentially, comprise Cell-APP. For our first application, we used Cell-APP to annotate training data for four adherent cell lines: HeLa, U2OS, HT1080, and RPE-1. To collect the inputs needed for Cell-APP (paired TL and nuclear fluorescence images), we imaged monocultures of each cell line in 96-well plates (see **Methods** for procedural details). We then processed images from each cell line separately using Cell-APP, resulting in four cell line-specific datasets (**Figure 2a**). Summary statistics on each dataset are available in **Table 2**. For training, we split each dataset into a training set (~70%) and a test set (~30%).

**Figure 2.**
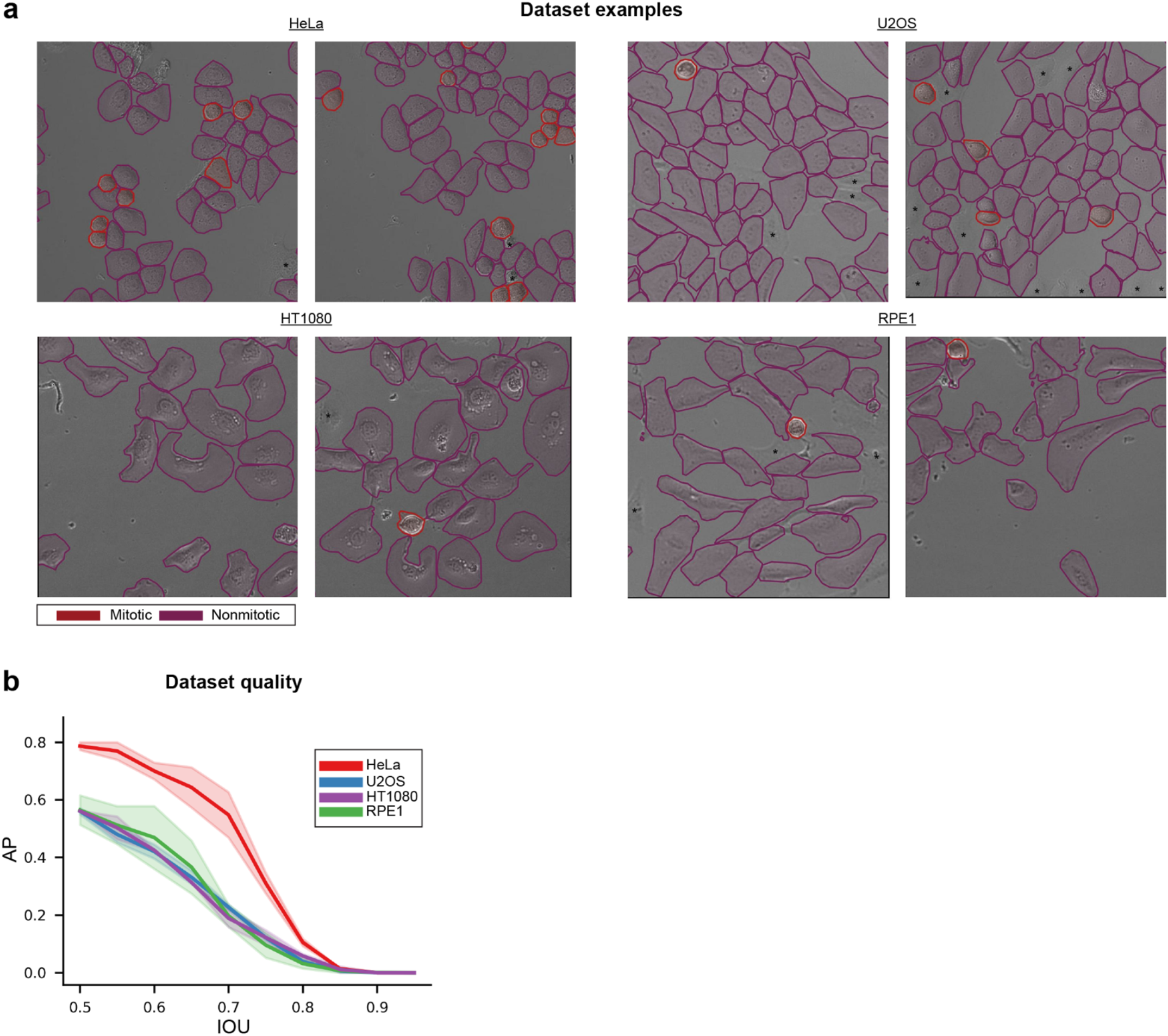
Cell-APP-generated datasets: examples and quality metrics. **a**, Example annotations from each of the four cell line-specific datasets, HeLa, U2OS, HT180, and RPE1. Asterisks denote cells missed by the pipeline. **b**, Comparison between hand annotations and Cell-APP-generated annotations for each cell line reveals that Cell-APP dataset quality differs across the cell line-specific datasets. AP was computed using IOU thresholds spanning 0.5 to 0.95 in increments of 0.05. Error bounds depict the mean ± standard error. Reasons for such quality differences are discussed in **Fig. S1**.

To assess the quality of the Cell-APP-annotated datasets, we hand-annotated one image from each training and test set (two per dataset) and computed average precisions (APs) between the Cell-APP annotations and hand-annotations (**Figure 2b**). AP at 50% intersection-over-union (IOU) exceeded 50% across all images, indicating acceptable performance. Based on these results and visual inspection, we concluded that the Cell-APP-annotated datasets were of sufficient quality to proceed with instance segmentation model training. We did, however, notice quality differences between the HeLa and U2OS, HT1080, and RPE-1 datasets (**Figure 2b**). These differences are attributable to errors in segmentation by µSAM. U2OS, HT1080, and RPE-1 cells have stellate or fusiform morphologies, unlike the predominantly cuboidal HeLa cells. On occasion, µSAM under-segments the outer edges of such stellate and fusiform cells (**Figure 2a**). µSAM also occasionally misidentifies these cells’ borders and over-segments them; this results in overlapping masks. Cell-APP removes one of any two overlapping masks (**Figure S1; Methods**). This preserves annotation integrity; however, it reduces dataset quality by causing datasets to contain unannotated cells (marked by asterisks in **Figure 2a**)

### Cell line-specific models

We next used these datasets to train cell-line-specific instance segmentation models. For this task, we used the model architectures and training framework offered by Detectron2, Meta AI’s Python library for object detection and segmentation (Yuxin Wu, 2019). To determine the optimal model architecture, we trained ResNet50, Vision Transformer-base (ViTB), and Vision Transformer-large (ViTL) backbone models on the HeLa training set. Based on mean average precision (mAP) and mean average recall (mAR) on the HeLa test set, we selected ViTL for subsequent applications (**Figure 3b; Methods**).

**Figure 3.**
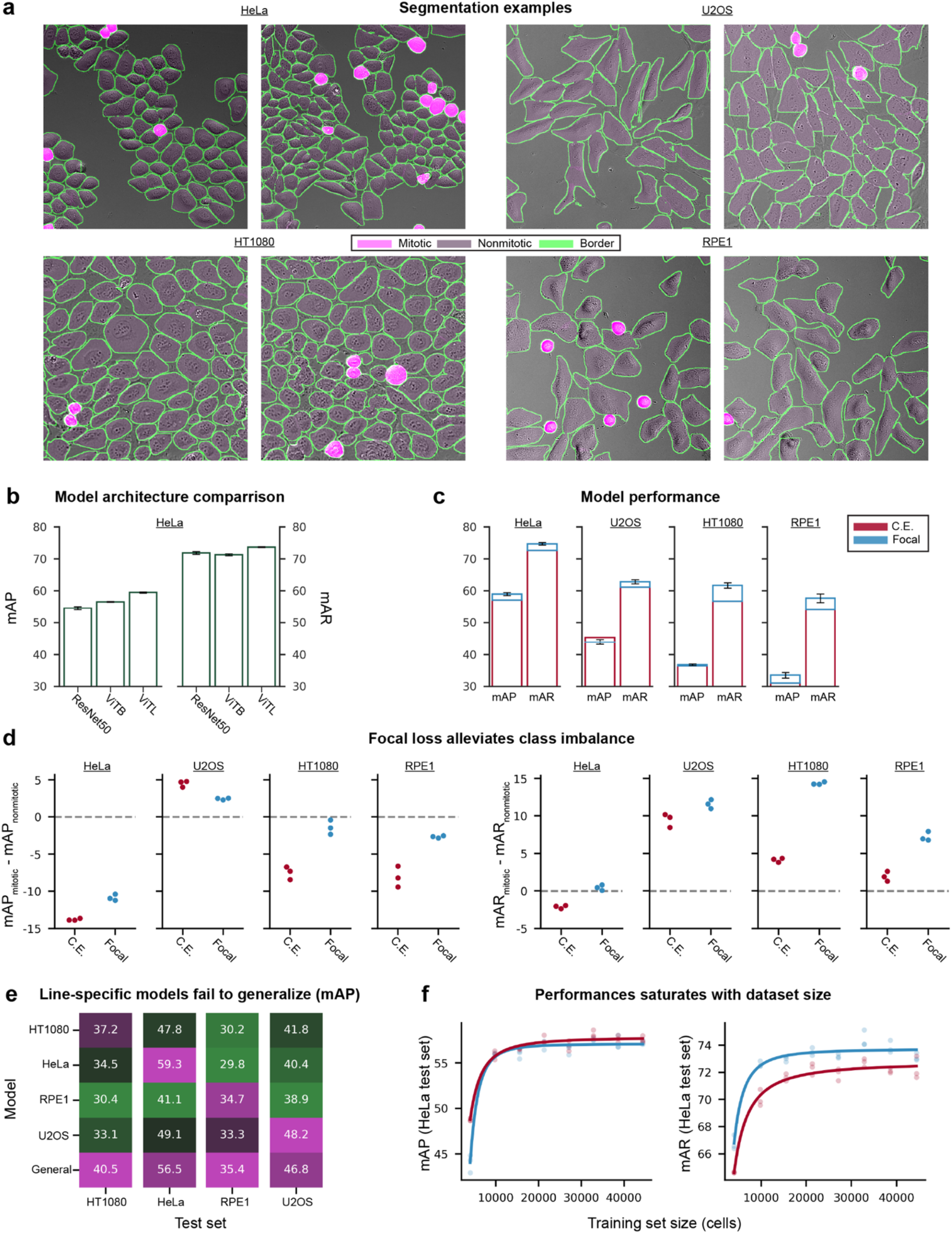
Cell-APP models accurately segment and classify cells. **a**, Segmentations and classifications generated by Cell-APP dataset trained models. <u>Cell line</u> denotes that the image displays <u>cell line</u> cells and was segmented by a model trained on the Cell-APP <u>cell line</u> dataset. **b**, To determine the ideal model architecture, Detectron2 models were trained on the HeLa dataset with varying backbones. Models were compared using mean average precision (mAP) and mean average recall (mAR), calculated as the respective means of AP and AR values across IOU thresholds spanning 0.5 to 0.95 in increments of 0.05. The ViTL backbone was chosen for subsequent experiments. **c**, Performance summary for cell line-specific models (models trained and evaluated on cell line-specific datasets). Blue top bars indicate the difference in performance between models trained with focal loss and models trained with cross-entropy (CE) loss. Stark differences in performance between cell line-specific models arise, in part, from differences in the quality of the testing dataset (**Fig. 2b, c). d**, Focal loss improves model performance on mitotic cells with respect to their performance on non-mitotic ones. The U2OS models is an exception. **e**, Cell line-specific models do not perform well on out-of-distribution (each other’s) testing datasets. However, a general model trained on ~11k cells of each line (*bottom row)* rivals in-distribution performance. **f**, Eight general models were trained on datasets of increasing size (from ~4k to ~45k cells) and equal proportion HeLa, U2OS, HT1080, and RPE1 cells. Evaluation of these models on the HeLa testing dataset shows that performance saturates after ~10k cells, suggesting a minimum sufficient dataset size. Similar results were found for the other three testing datasets (**Fig. S3**). All error bars depict mean ± standard error (performance evaluated at *n = 3* epochs, showing model convergence). We fit Hill functions to the data for visualization purposes only.

We trained three additional ViTL backbone models—one per cell line. Segmentations produced by each cell line-specific model, as well as the mAP and mAR between each model and its respective test set, are shown in **Figures 3a and 3c** (**red bars**). It is important to note that the differences in these metrics across models are not entirely attributable to differences in model performance. Unannotated cells in the test set (which result from µSAM segmentation errors as discussed above) will result in correct model predictions being classified as false positives, thus causing mAP to decrease (**Methods**). The extent to which these errors occur differs across test sets (**Figure 2b**). To better understand model performance, we computed APs between our models’ predictions and hand-annotations. The results (**Figure S2c**) confirmed that differences in model performance are less drastic than **Figure 3c** suggests.

### Focal loss for addressing class imbalance

Despite overall high performance, our models showed a consistent weakness: they were less precise in segmenting mitotic cells than non-mitotic ones (**red circles in Figure 3d**). The U2OS model was an exception to this trend for reasons that remain unclear. This precision imbalance may stem from class imbalance in our training datasets; the cell line-specific datasets have a 20:1 ratio of non-mitotic to mitotic annotations on average (**Table 2**). This class imbalance is biological, reflecting the fact that cells spend much less time in mitosis than not. For example, HeLa cells spend ~30 minutes in mitosis during their ~18-hour cell cycle. Hence, non-mitotic cells will naturally outnumber mitotic ones in an unsynchronized cell population. We partially mitigated this imbalance by using mitotic poisons to increase our cells’ time-in-mitosis (we treated cells in the training sets only; **Methods**).

To address this weakness, we retrained each model using categorical focal loss instead of categorical cross-entropy loss. Focal loss is a variant of cross-entropy that biases models to learn from “hard” (low-confidence) predictions by downweighting the loss from “easy” (high-confidence) predictions (Lin *et al*., 2017). We find that focal loss increases: (1) overall mAP and mAR, (2) mitotic-mAP relative to non-mitotic-mAP, and (3) mitotic-mAR relative to non-mitotic-mAR (**blue bars and circles in Figures 3c-d**). The exception to this trend is, again, the U2OS model; focal loss decreased its overall mAP and mitotic-mAP relative to non-mitotic mAP (**Figure 3c-d**). We note that none of these trends depend on model architecture (**Figure S3**).

In agreement with this precision imbalance, we found that our models’ most common error is misclassifying non-mitotic cells as mitotic (**Figure S5**). It is important to note that the models make these misclassifications with low (~5%) confidence; therefore, they can be filtered out (**Figure S5; Methods**). Moreover, these misclassifications are “double predictions” (i.e., the second mask predicted for a given cell); therefore, removing them does not result in unsegmented cells (**Figure S5**). These double predictions arise because we permit up to 2,000 mask predictions per image to avoid under-segmentation.

One possible explanation for this error mode and the resulting precision imbalance is that the models confuse non-mitotic cells with prophase cells, which are labeled as mitotic in the training data. A model-segmented and classified prophase cell is shown in **Figure 4a**. Prophase cells often resemble non-mitotic cells in both size and shape (**Figure S1b**). This resemblance is pronounced in cuboidal cell lines, like HeLa, which lack stellate or fusiform extensions that would otherwise retract during pre-mitotic rounding. Hence, this confusion may also explain why the HeLa model has the greatest non-mitotic/mitotic precision imbalance (**Figure 3d**), despite its training set having the greatest mitotic-to-non-mitotic ratio (**Table 2**).

**Figure 4.**
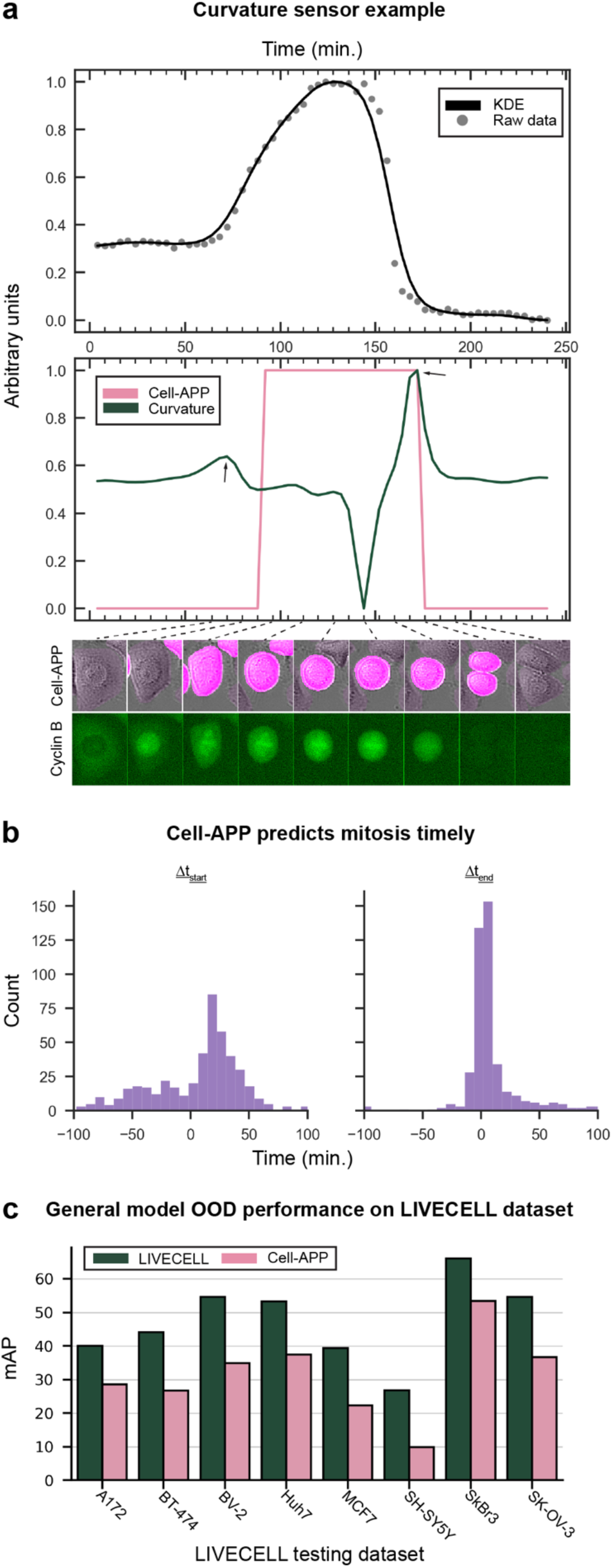
Cyclin B validates model classifications. LIVECELL validates model segmentations. **a**, *Top*, Cyclin B trace and kernel density estimation. *Middle*, curvature of the same Cyclin B trace and Cell-APP’s classification of the corresponding cell. Local maxima of the curvature (see arrows) are biological estimates of mitotic start and stop. Cell-APP = 1 indicates a mitotic prediction for that time point. *Bottom*, Cell-APP prediction and Cyclin B montages; dark pink indicates a Cell-APP mitosis prediction; dashed lines point to the shown time point. **b**, *Left*, Cell-APP mitotic start predictions lag the biological estimate with a median of 16 ± 46.5 minutes. *Right* Cell-APP mitotic stop predictions lag the biological estimate with a median of 4 minutes ± 14.9 minutes (*n = 488*). **c**, Performance of the Cell-APP general model (training dataset of ~45k cells) is comparable to the performance of the LIVECell general model (training dataset of ~one million cells), despite having never seen instances of these eight cell lines. mAP was computed as the mean of AP values spanning IOU thresholds from 0.5 to 0.95 in increments of 0.05.

### General models trained on multi-cell line datasets

A central goal in cell segmentation is to produce a single model that performs well across diverse cell lines. To assess this ability in our models, we evaluated each cell line-specific model on the three other “non-target” test sets (where “non-target” test sets are ones containing cell lines the model did not train on). The performance of these models was, in each case, worse than the performance of the target set-trained model (**Figure 3e**).

Previous work has shown that training a model on multiple cell lines does not compromise its performance on individual cell lines (Edlund *et al*., 2021). Motivated by these findings, we set out to construct a multi-cell line dataset and train a single model capable of performing well across all cell line-specific test sets. To create the multi-cell line dataset, we collated images and annotations from the HeLa, U2OS, HT1080, and RPE-1 training datasets. In doing this, we used stratified sampling to ensure that each cell line and class was uniformly represented in the multi-cell line dataset (**Table 3**). This dataset, which we call the “general dataset”, contains ~11,000 annotations from each cell line, ~2,000 being mitotic and ~42,000 being non-mitotic. We additionally used this procedure to create seven more, uniformly distributed, general datasets of increasing size; the smallest containing ~4000 annotations, ~200 being mitotic, and ~3,400 being non-mitotic.

With these general datasets in hand, we trained a model that generalizes across cell line-specific test sets. To do this, we trained two ViTL-backbone models on the largest (~44,000 cells) general dataset—one using cross-entropy loss and one using focal loss. We evaluated these general models on the individual HeLa, U2OS, HT1080, and RPE-1 test sets. In all cases, the general models outperformed the cell line-specific models that were not trained on the target set, and even the target set-trained model in some cases (**Figure 3e**). We report mAP values from the focal loss-trained general model in **Figure 3e**; however, the same trends hold for the model trained using cross-entropy loss.

Having established that Cell-APP can be used to train a general model that performs across cell lines, we next asked: how much data is required to achieve this level of performance? To answer this, we trained ViTL-backbone models on each of the eight, increasingly large general datasets. We trained two models on each dataset—one using cross-entropy loss and one using focal loss. We found that across all test sets, mAP and mAR begin to plateau once the training set exceeds 10,000 cells (**Figure 3f; Figure S3c**). Focal loss consistently improved mAR but not mAP (**Figure 3f; Figure S3c**). These results show that a relatively modest number of annotated examples is sufficient to train a generalist cell segmentation model. We note, though, that annotating even 10,000 cells is prohibitively tedious to do by hand.

Finally, as a benchmark, we evaluated our focal loss-trained general model on external, cell line-specific datasets from LIVECell (**Figure 4c**). Our general model was not trained on any of the cell lines in these datasets. Based on mAP, our model’s performance is comparable to, but worse than, that of the LIVECell generalist model. This is expected; LIVECell’s generalist was trained to segment the cell lines in these datasets, and it was trained on a dataset containing over one million cells (Edlund *et al*., 2021).

### Biologically validating Cell-APP model classifications

We finally sought to assess the accuracy of our models’ cell-cycle classifications in a “real-life” experiment. To do this, we compared model classifications on time-lapse movies to a biological marker of mitotic timing: Cyclin B abundance. Cyclin B is an essential mitotic protein; it starts accumulating in late G2 and is degraded in metaphase and anaphase, with the end of its degradation marking the end of mitosis (Gavet and Pines, 2010a, b). Therefore, to capture these dynamics, we imaged genome-edited HeLa cells, expressing mNeonGreen-Cyclin B1 for upwards of 24 hours in 4-minute intervals. Visual inspection revealed that the onset of Cyclin B accumulation (G2) and the end of its degradation (mitosis end) coincide with sharp bends in the graph of Cyclin B abundance vs. time (**Figure 4a**). Precisely, these bends are the local maxima of the graph’s curvature. We, therefore, used the positions of these two local maxima as indicators of mitotic start and stop time.

We segmented and classified each frame of the time-lapse movies using the focal loss-trained HeLa model, and recorded Cyclin B concentration as the mean fluorescence within individual Cell-APP segmentations. From these frame-by-frame classifications, we extracted the first and last time points in which Cell-APP called a given cell “mitotic”; we refer to these as the Cell-APP estimates of mitotic start and stop time (**Methods**). We used simple differences to compare the Cell-APP and biological estimates (**Methods**). Reporting on the median ± standard deviation, we found that Cell-APP predicts mitotic start 16 ± 46.5 minutes after Cyclin B and mitotic end 4 ± 14.9 minutes after Cyclin B (**Figure 4b**). The discrepancy between Cell-APP’s and Cyclin B’s estimation of mitotic start arises from the fact that Cyclin B’s estimation more faithfully indicates the beginning of late G2, not mitosis. In other words, Cyclin B’s estimation is inherently early, and Cell-APP’s prediction of mitotic start, which relies on distinct concomitant changes in cell morphology, is likely more accurate. From these results, we conclude that Cell-APP classifications are temporally accurate (Gavet and Pines, 2010a).

## Discussion

Cell-APP is a general-purpose method for annotating cells in microscopy images and thereby creating instance segmentation training data. Our applications of Cell-APP have validated the method by yielding segmentation models that achieve an average AP50 of ~80 on hand-annotated data (**Figure S2b**). These models are provided as ready-to-use tools for the cell biology community. The method largely excises the need for hand-annotation from cell segmentation model training pipelines and serves as an example of how creatively using current technologies (e.g., SAM) enables the development of novel image analysis tools in biology (Dixon *et al*., 2025; Franco-Barranco *et al*., 2025).

We intend for Cell-APP to be used in two ways. First, we hope researchers will employ Cell-APP to train cell segmentation models, customized to their cell lines and imaging setup. These models will automate TL microscopy analysis, thus enabling high-throughput studies and accelerating biological discovery. Second, we intend to create a repository where users can upload their Cell-APP-generated datasets. By training deep learning models on this continuously growing collection of data, we hope to create a truly general, TL cell segmentation tool. The promise of this second use case is supported by the performance of our “general model.”

There are many future directions for Cell-APP. First, the current classification pipeline only distinguishes between mitotic and non-mitotic nuclei. In select cases, we can identify anaphase nuclei by re-clustering the mitotic cluster; however, the identified anaphase clusters tend to be contaminated with dead and metaphase cells (**Figure S1b**). By modifying the pipeline to use geometric features across multiple spatial scales (as opposed to one), or by implementing a more sophisticated re-clustering protocol, it may reliably distinguish between prophase, metaphase, anaphase, and dead cells (Berg *et al*., 2019). Second, we have only used the classification pipeline to create cell-cycle labels. In principle, it could create classifications from other fluorescence signals. For example, it could distinguish between populations of cells with differing gene expression profiles (by visualizing transcription factor concentrations) or differing morphologies (by visualizing the cell membrane). Implementing such flexibility would increase Cell-APP’s utility. We caution, however, that some fluorescence-based classifications may not correspond to discernible features in TL images. In this case, the trained cell segmentation models may not learn to make accurate classifications. Last, Cell-APP mask generation is only as good as SAM (or µSAM). As these models improve, or as more advanced promptable segmentation models emerge, Cell-APP’s utility will likewise improve.

## Methods

### Time-lapse live-cell imaging

To collect images used for Cell-APP dataset annotation, cells were imaged using an ImageXpress Nano Automated Imaging System (Molecular Devices) equipped with a SOLA Light Engine (Lumencore) as the excitation source, and a 20X, 0.46 NA objective. Images in the Cell-APP datasets are of cells seeded in 96-well plates and imaged using a 20X, 0.46 NA objective. During imaging, we supplied humidified 5% CO2 to the environment chamber; this chamber was kept at 37 °C. To facilitate nucleus visualization, we used cell lines stably expressing Histone 2B-mCherry.

The following protocol was used for image collection: 24 hours before imaging, cells stably expressing Histone 2B-mCherry were seeded at varying confluency in a 96-well plate. Cells were seeded in sets of three wells; images from two wells were used for training data, and images from the third well were used for testing. Thirty minutes before imaging, the training data wells were treated with 12.5-50 nM GSK923295 to increase the number of mitotic cells imaged (Bennett *et al*., 2015). Increasing the number of mitotic cells aided our classification procedure and reduced dataset class imbalance. We then imaged each well for 6-8 hours in 2-hour intervals. During each interval, images were captured in pairs (transmitted-light and nuclear fluorescence) at three focal planes: one “in-focus”, one “above-focus”, and one “below-focus.” To clarify, focus was only adjusted on the transmitted-light images, the nuclear fluorescence images remained “in-focus.”

### Mask annotation with SAM

The pipeline is written in Python 3.11 and uses NumPy and Sci-Kit Image extensively (van der Walt *et al*., 2014; Harris *et al*., 2020). To compute centroids and bounding boxes for each nucleus, we implemented a pre-processing and thresholding procedure. The procedure first smooths the nuclear fluorescence image using Gaussian convolution. It then subtracts the background using a morphological white top-hat operation. Both operations prepare the image for thresholding. It then applies erosion to ensure that nuclei are not touching; this is necessary to distinguish between neighboring nuclei. It then thresholds the image and labels individual nuclei. via connected component labeling. The result of this process is one mask per nucleus. It is then straightforward to compute each mask’s centroid. A centroid and a bounding box centered on said centroid are then computed for each mask. The size of the bounding box is determined by the size of the mask.

Users can control many parameters in this process, including the Gaussian kernel, the white top-hat kernel, the erosion kernel, and the thresholding algorithm. We provide the parameters used for HeLa, U2OS, HT1080, and RPE1 dataset creation here: https://github.com/anishjv/cell-AAP/tree/main/notebooks. Of note: bounding boxes that extend beyond the image are discarded. This is done for two reasons: (1) the same bounding boxes are used to crop out cells for classification. If extra-image bounding boxes are not discarded, crops may be of drastically different sizes, which will bias classification. (2) SAM will not process bounding boxes that are not fully contained within the image.

### Classification via unsupervised learning

The features used for annotation classification were constant across datasets; a complete list is provided in **Table 1**. These features are adjustable pipeline parameters; users can remove features and add new ones, so long as they are compatible with the skimage function “regionprops().” Parameter values used for both Uniform Manifold Approximation and Projection (UMAP) and Hierarchical Density-based Spatial Clustering of Applications with Noise (HDBSCAN) can be found at https://github.com/anishjv/cell-AAP/tree/main/notebooks (Campello *et al*., 2013; McInnes *et al*., 2018).

In practice, we found that our pipeline works best on datasets with over 1,000 cells of each class. With fewer cells, two errors may occur: (1) UMAP may be biased by intra-class heterogeneity and produce extra, spurious classes, or (2) if one class is significantly more abundant than the other (ratio greater than 10:1), the pipeline may find only one cluster. The first error is easily corrected by visually inspecting and merging classes; we recommend collecting additional, balanced data to correct the second.

### Hand-annotations

Two images from each of the HeLa, U2OS, HT1080, and RPE1 datasets (one from the training set and one from the test set) were hand-annotated to benchmark the pipeline’s performance. Cells in these images were annotated as polygons by two independent annotators, using both the transmitted light and nuclear fluorescence images. Both annotators used https://www.makesense.ai/. Using both the transmitted light and nuclear fluorescence images when annotating. Using the nuclear fluorescence image allowed annotators to localize each cell and put them on an even playing field with the Segment Anything Model (SAM); SAM is informed of the cell’s location before segmenting.

To benchmark the pipeline’s performance, we computed the average precision (AP) between the pipeline’s annotations and our hand annotations. We followed the Common Objects in Context (COCO) evaluation standards, computing AP at intersection-over-union (IOU) thresholds spanning 0.5 to 0.95 in increments of 0.5 (Lin *et al*., 2014). We additionally used the test set-originating hand-annotated images to benchmark model performance. In doing so, we again followed the COCO evaluation standards.

### Model architecture

We used Detectron2, Meta AI’s object-detection platform, to develop all segmentation models (He *et al*., 2017). Detectron2 models are written and optimized with PyTorch (Paszke *et al*., 2019). To determine a model architecture for all future experiments, we trained three Mask-Region-based Convolutional Neural Networks (Mask-RCNN) on the Cell-APP HeLa dataset. The three models differed in backbone, using ResNet50, Vision Transformer Base (ViTB), and Vision Transformer Large (ViTL), respectively (He *et al*., 2016; Dosovitskiy *et al*., 2020). We evaluated these models via AP and average recall (AR), on the HeLa test set.

### Model training

For all models, we started training from weights learned on ImageNet-1K. Training was conducted on one NVIDIA A40 GPU with 48 GB of RAM. We trained each model for 2000 epochs, stored weights every 200 epochs, and evaluated the model on the test set every 100 epochs to probe for overfitting. We employed a pseudo early stopping protocol, in which we allowed models to train for the full 2000 epochs, but used the set of weights from the highest mAP epoch for post-training experiments. We used Adam with weight decay (AdamW) as our model optimizer with β_1_ = 0.9 and β_2_ = 0.99 (Adam, 2014). We used a stepwise learning rate decay and a 250-iteration-long linear learning rate warm-up. Models were trained on 1024×1024 pixel images. The patch size for the vision-transformer models was set at 16. All training used multi-scale and intensity-based data augmentations, including cropping, resizing, random contrast, and brightness adjustments. We provide model configurations here: https://zenodo.org/communities/cellapp.

### Focal loss for addressing class imbalance

Binary cross-entropy is a canonical loss function for training deep learning models to classify objects as one of two classes. It is defined as:

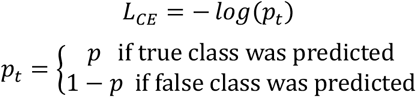

Here *p* is the model’s prediction for the probability that the predicted class is correct; *p*_*t*_ is the model’s prediction for the probability that the true class is correct. During training, this loss function is computed for each instance. The mean across instances is the model’s total class loss.

Focal loss biases models to learn “hard” (low *p*_*t*_) instances by downweighting the loss of “easy” (high *p*_*t*_) instances (Lin *et al*., 2017). Explicitly, focal loss is defined as:

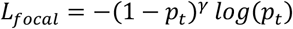

Here, the scalar multiplier (1 − *p*_*t*_)^*γ*^ causes high *p*_*t*_ instances to have minimal loss compared to low p_*t*_ instances. The focusing parameter, *γ*, is defined to be greater than zero and controls the extent of weighting. For model training, we used γ = 2. An exploration on the effect of focusing parameter value on model performance is provided in **Figure S3**.

### Evaluation metrics

We chose mean AP (mAP) and mean AR (mAR) as our primary evaluation metrics. These metrics are defined by three more fundamental metrics: intersection-over-union (IOU), precision, and recall. Precision and recall are defined as:

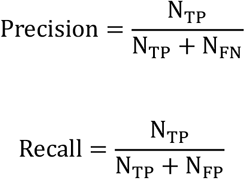

TP, FP, and FN are classifications assigned to the segmentations that our models predict. They denote whether a segmentation matches the ground-truth (true positive, TP), does not match the ground truth (false positive, FP), or if the model failed to predict an instance when there should have been one (false negative, FN). *N*_*x*_ denotes the number of annotations classified as *x*.

IOU is used to determine whether or not an instance “matches” the ground-truth:

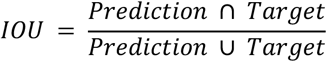

IOU measures the overlap between predicted instances (such as segmentation or bounding boxes) and the nearest ground-truth instances. In practice, an IOU threshold *t*_*IOU*_ is chosen such that instances with IOU > *t*_*IOU*_ are classified as TP and instances with IOU < *t*_*IOU*_ are classified as FP.

To compute AP, first, all predicted segmentations are separated by class and sorted in descending order by model confidence. Then, for one class, precision and recall are computed using only the two lowest confidence segmentations. Precision-recall pairs continue to be computed, adding the remaining lowest confidence segmentation to the computation pool each time. This process is repeated until all segmentations have been included in the computation. The resulting precision-recall pairs form a discrete function *p*_*c*_(*r*). The integral of this function is the class-wise AP.

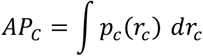

This process is repeated for the other classes, and the mean of all class-wise AP metrics is AP.

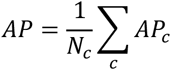

Here *N*_*c*_ is the number of classes. It is critical to remember that AP depends on the IOU threshold used, as this threshold defines TP and FP. mAP is defined as the mean of AP metrics computed with different IOU thresholds. Explicitly, COCO has chosen the IOU values spanning 0.5 to 0.95 in increments of 0.05. We adopted this definition for our reporting.

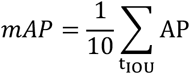

mAR is computed similarly. The functional relationship between precision-recall pairs is simply reversed.

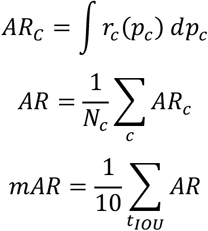

### Metric reporting

In computing mAP and mAR, we chiefly followed COCO guidelines. However, we adjusted maxDets, the parameter controlling the maximum number of model-predicted segmentations, from 100 to 2000. This was necessary as Cell-APP datasets often contain images with over 1000 cell annotations. We did not post-process model outputs when computing mAP and mAR. When reporting metrics, such as mAP and mAR, a set of model weights must be chosen for inference. For all training experiments (shown in **Figure 3b-d** and **Figure 3f**), we computed metrics using weights from the three epochs with the highest mAP. We report three values to show the convergence of our models’ performance. For the data in **Figure 2c**, we report metrics from the three highest mAP epochs of the focal loss models.

For the cross-cell line experiments, we chose cell line-specific model weights using the pseudo early stopping protocol mentioned in **Model training**. In rows 1-4 of **Figure 3e**, we report metrics from focal loss-trained HeLa, HT1080, and RPE1 models, and a cross-entropy-trained U2OS model. For the general models, we used the set of weights from the large general model (training set size ~ 45k cells), which performed best across the HeLa, U2OS, RPE1, and HT1080 test sets. The evaluation of these models on each test set occurred after model training. We report one mAP value for each model-test set pair, which compares that model’s predictions against that test set’s annotations. We used the same set of weights for the general model in our comparison against LIVECELL (**Figure 4c**). It is important to note that these metrics are comparable to, but do not match, those found in **Table 4**. All metrics found in **Table 4** were computed using weights from the three epochs with the highest mAP.

### Cyclin B microscopy analysis

Genome-edited HeLa cells expressing mNeonGreen-CyclinB1 were seeded in 96-well plates. We imaged these cells for upwards of 24 hours in 4-minute intervals using the previously described ImageXpress Nano microscopy setup. We also imaged one well containing fluorescent media (Dulbecco’s Modified Essential Medium (DMEM)) and another containing Fluorobrite DMEM. These images were used for illumination correction and background subtraction, respectively.

We used the Cell-APP HeLa model to identify and classify cells in these images. We used the Python program, TrackPy, for cell tracking (Allan, 2021).

### Segmentation post-processing

In two ways, we processed segmentations produced by the Cell-APP HeLa model before tracking them. First, we removed any segmentations that the model made with less than 25% confidence. As mentioned in **Results**, such segmentations are typically redundant and may be misclassifications. Second, if any two segmentations overlap with more than 50% IOU, we remove the lower confidence one. Again, the greater than 50% IOU suggests that the removed segmentation was the second segmentation for that cell (i.e., it was redundant). Our models make redundant segmentations because we allow them to make up to 2,000 predictions. Our rationale for such a high limit is that it is easier to remove redundant segmentations than to estimate the number of cells in a TL image.

### Curvature sensor for predicting mitosis

To quantify a given cell’s intracellular Cyclin B abundance, we computed the mean intensity within the corresponding Cell-APP segmentation. Concentration quantification and tracking allowed us to construct discrete traces of the form [*CyclinB*](*t*) for each cell; *t* is time. We used the curvature, *κ*(*t*), of our CyclinB 15 traces to biologically estimate the timepoints at which each cell entered and exited mitosis. Since differentiation drastically reduces signal-to-noise ratio, we smoothed each trace using kernel density estimation before computing *κ*(*t*).

The curvature of a curve describes how quickly a line tangent to the curve changes direction as it is moved along the curve. We can derive the curvature of a trace by considering a parametric plane curve,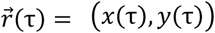. We will call our parameter τ to distinguish it from time. Every plane curve can be reparameterized by its arclength. Arclength is defined as:

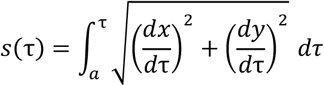

When parameterized in terms of arclength, the curve takes the form: 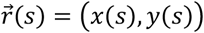. Now each step along the curve is equivalent to traveling one unit of distance. The derivative of 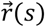 is called the unit tangent function,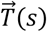. It defines a line of unit length tangent to every point, *s*, on the curve. The rate at which this function changes direction, 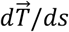, is the curvature.

After doing the algebra that corresponds to this description, we find that:

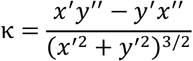

Here, prime notation signifies the derivative with respect to the parameter τ. We also note that a trace is just a special case of a plane curve in which *x*(τ) = *t* and *y*(*τ*) = *y*(*t*). In this case, the curvature reduces to:

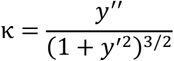

The timepoints of the two most prominent local extrema of κ were taken as biological estimates of mitotic start and stop. These were found using Sci-Kit Learn’s neighborhood value comparison algorithm. We define Δ*t*_*start*_ and Δ*t*_*stop*_ as:

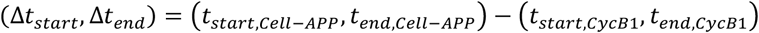

Physically, Δ*t*_*start*_ is the delay in Cell-APP’s prediction of mitosis start and Δ*t*_*stop*_ is the delay in Cell-APP’s prediction of mitosis end.

## Acknowledgments

We thank Dr. Matthew O’Meara and Dr. Idse Heemskerk for reading drafts of this manuscript and providing insightful suggestions. AJV thanks Mishan Gagnon for always enjoyable and insightful conversations, including the one that introduced AJV to SAM. AJV also thanks Jennifer Guan’ for hand-annotating microscopy images and the entire Joglekar Lab for their support. This work was supported by R35-GM126983 to APJ for NIGMS.

## References

Adam, K.D.B.J. (2014). A method for stochastic optimization. arXiv preprint arXiv:1412.6980 1412.

Allan, D.B.C., Thomas A. Keim, Nathan C. van der Wel, Casper M. (2021). A method for stochastic optimization. 2025: Zenodo.

Archit, A., Freckmann, L., Nair, S., Khalid, N., Hilt, P., Rajashekar, V., Freitag, M., Teuber, C., Spitzner, M., Tapia Contreras, C., Buckley, G., von Haaren, S., Gupta, S., Grade, M., Wirth, M., Schneider, G., Dengel, A., Ahmed, S., and Pape, C. (2025). A method for stochastic optimization. Nat Methods 22, 579–591.

Bennett, A., Bechi, B., Tighe, A., Thompson, S., Procter, D.J., and Taylor, S.S. (2015). A method for stochastic optimization. Oncotarget 6, 20921–20932.

Berg, S., Kutra, D., Kroeger, T., Straehle, C.N., Kausler, B.X., Haubold, C., Schiegg, M., Ales, J., Beier, T., and Rudy, M. (2019). A method for stochastic optimization. Nature methods 16, 1226–1232.

Brown, T., Mann, B., Ryder, N., Subbiah, M., Kaplan, J.D., Dhariwal, P., Neelakantan, A., Shyam, P., Sastry, G., and Askell, A. (2020). A method for stochastic optimization. Advances in neural information processing systems 33, 1877–1901.

Campello, R.J., Moulavi, D., and Sander, J. (2013). A method for stochastic optimization. Pacific-Asia conference on knowledge discovery and data mining, 160–172.

Carpenter, A.E., Jones, T.R., Lamprecht, M.R., Clarke, C., Kang, I.H., Friman, O., Guertin, D.A., Chang, J.H., Lindquist, R.A., and Moffat, J. (2006). A method for stochastic optimization. Genome biology 7, R100.

Clute, P., and Pines, J. (1999). A method for stochastic optimization. Nature cell biology 1, 82–87.

Cohen, E., and Uhlmann, V. (2021). A method for stochastic optimization. 2021 IEEE 18th International Symposium on Biomedical Imaging (ISBI), 640–644.

Dixon, J.C., Frick, C.L., Leveille, C.L., Garrison, P., Lee, P.A., Mogre, S.S., Morris, B., Nivedita, N., Vasan, R., and Chen, J. (2025). A method for stochastic optimization. Cell Systems 16.

Dosovitskiy, A., Beyer, L., Kolesnikov, A., Weissenborn, D., Zhai, X., Unterthiner, T., Dehghani, M., Minderer, M., Heigold, G., and Gelly, S. (2020). A method for stochastic optimization. arXiv preprint arXiv:2010.11929.

Edlund, C., Jackson, T.R., Khalid, N., Bevan, N., Dale, T., Dengel, A., Ahmed, S., Trygg, J., and Sjögren, R. (2021). A method for stochastic optimization. Nature methods 18, 1038–1045.

Falk, T., Mai, D., Bensch, R., Çiçek, Ö., Abdulkadir, A., Marrakchi, Y., Böhm, A., Deubner, J., Jäckel, Z., and Seiwald, K. (2019). A method for stochastic optimization. Nature methods 16, 67–70.

Fazeli, E., Roy, N.H., Follain, G., Laine, R.F., von Chamier, L., Hänninen, P.E., Eriksson, J.E., Tinevez, J.-Y., and Jacquemet, G. (2020). A method for stochastic optimization. F1000Research 9, 1279.

Fishman, D., Salumaa, S.O., Majoral, D., Laasfeld, T., Peel, S., Wildenhain, J., Schreiner, A., Palo, K., and Parts, L. (2021). A method for stochastic optimization. Journal of Microscopy 284, 12–24.

Franco-Barranco, D., Andrés-San Román, J.A., Hidalgo-Cenalmor, I., Backová, L., González-Marfil, A., Caporal, C., Chessel, A., Gómez-Gálvez, P., Escudero, L.M., Wei, D., Muñoz-Barrutia, A., and Arganda-Carreras, I. (2025). A method for stochastic optimization. Nature Methods 22, 1124–1126.

Gavet, O., and Pines, J. (2010a). Activation of cyclin B1–Cdk1 synchronizes events in the nucleus and the cytoplasm at mitosis. Journal of Cell Biology 189, 247–259.

Gavet, O., and Pines, J. (2010b). Progressive activation of CyclinB1-Cdk1 coordinates entry to mitosis. Developmental cell 18, 533–543.

Harris, C.R., Millman, K.J., van der Walt, S.J., Gommers, R., Virtanen, P., Cournapeau, D., Wieser, E., Taylor, J., Berg, S., Smith, N.J., Kern, R., Picus, M., Hoyer, S., van Kerkwijk, M.H., Brett, M., Haldane, A., Del Rio, J.F., Wiebe, M., Peterson, P., Gerard-Marchant, P., Sheppard, K., Reddy, T., Weckesser, W., Abbasi, H., Gohlke, C., and Oliphant, T.E. (2020). A method for stochastic optimization. Nature 585, 357–362.

He, K., Gkioxari, G., Dollár, P., and Girshick, R. (2017). A method for stochastic optimization. Proceedings of the IEEE international conference on computer vision, 2961–2969.

He, K., Zhang, X., Ren, S., and Sun, J. (2016). A method for stochastic optimization. Proceedings of the IEEE conference on computer vision and pattern recognition, 770–778.

Kirillov, A., Mintun, E., Ravi, N., Mao, H., Rolland, C., Gustafson, L., Xiao, T., Whitehead, S., Berg, A.C., and Lo, W.-Y. (2023). A method for stochastic optimization. Proceedings of the IEEE/CVF international conference on computer vision, 4015–4026.

Lin, T.-Y., Goyal, P., Girshick, R., He, K., and Dollár, P. (2017). A method for stochastic optimization. Proceedings of the IEEE international conference on computer vision, 2980–2988.

Lin, T.-Y., Maire, M., Belongie, S., Hays, J., Perona, P., Ramanan, D., Dollár, P., and Zitnick, C.L. (2014). A method for stochastic optimization. European conference on computer vision, 740–755.

Ling, C., Majurski, M., Halter, M., Stinson, J., Plant, A., and Chalfoun, J. (2020). A method for stochastic optimization. Proceedings of the IEEE/CVF Conference on Computer Vision and Pattern Recognition Workshops, 966–967.

McInnes, L., Healy, J., and Melville, J. (2018). A method for stochastic optimization. arXiv preprint arXiv:1802.03426.

Paszke, A., Gross, S., Massa, F., Lerer, A., Bradbury, J., Chanan, G., Killeen, T., Lin, Z., Gimelshein, N., and Antiga, L. (2019). A method for stochastic optimization. Advances in neural information processing systems 32.

Patel, G., Tekchandani, H., and Verma, S. (2019). A method for stochastic optimization. 2019 International Conference on Advances in Computing, Communication and Control (ICAC3), 1–5.

Rivenson, Y., Liu, T., Wei, Z., Zhang, Y., De Haan, K., and Ozcan, A. (2019). A method for stochastic optimization. Light: Science & Applications 8, 23.

Schmidt, U., Weigert, M., Broaddus, C., and Myers, G. (2018). A method for stochastic optimization. International conference on medical image computing and computer-assisted intervention, 265–273.

Schwendy, M., Unger, R.E., and Parekh, S.H. (2020). A method for stochastic optimization. Bioinformatics 36, 3863–3870.

Stringer, C., and Pachitariu, M. (2025). A method for stochastic optimization. Nature methods 22, 592–599.

Stringer, C., Wang, T., Michaelos, M., and Pachitariu, M. (2021). A method for stochastic optimization. Nature methods 18, 100–106.

Ulicna, K., Vallardi, G., Charras, G., and Lowe, A.R. (2021). A method for stochastic optimization. Frontiers in Computer Science 3, 734559.

van der Walt, S., Schonberger, J.L., Nunez-Iglesias, J., Boulogne, F., Warner, J.D., Yager, N., Gouillart, E., Yu, T., and scikit-image, c. (2014). A method for stochastic optimization. PeerJ 2, e453.

Yuxin Wu, A.K., Francisco Massa, Wan-Yen Lo, Ross Girshick. (2019). Detectron2, https://github.com/facebookresearch/detectron2.

