## Supplemental materials for "Cell-APP: A generalizable method for cell annotation and cell-segmentation model training"

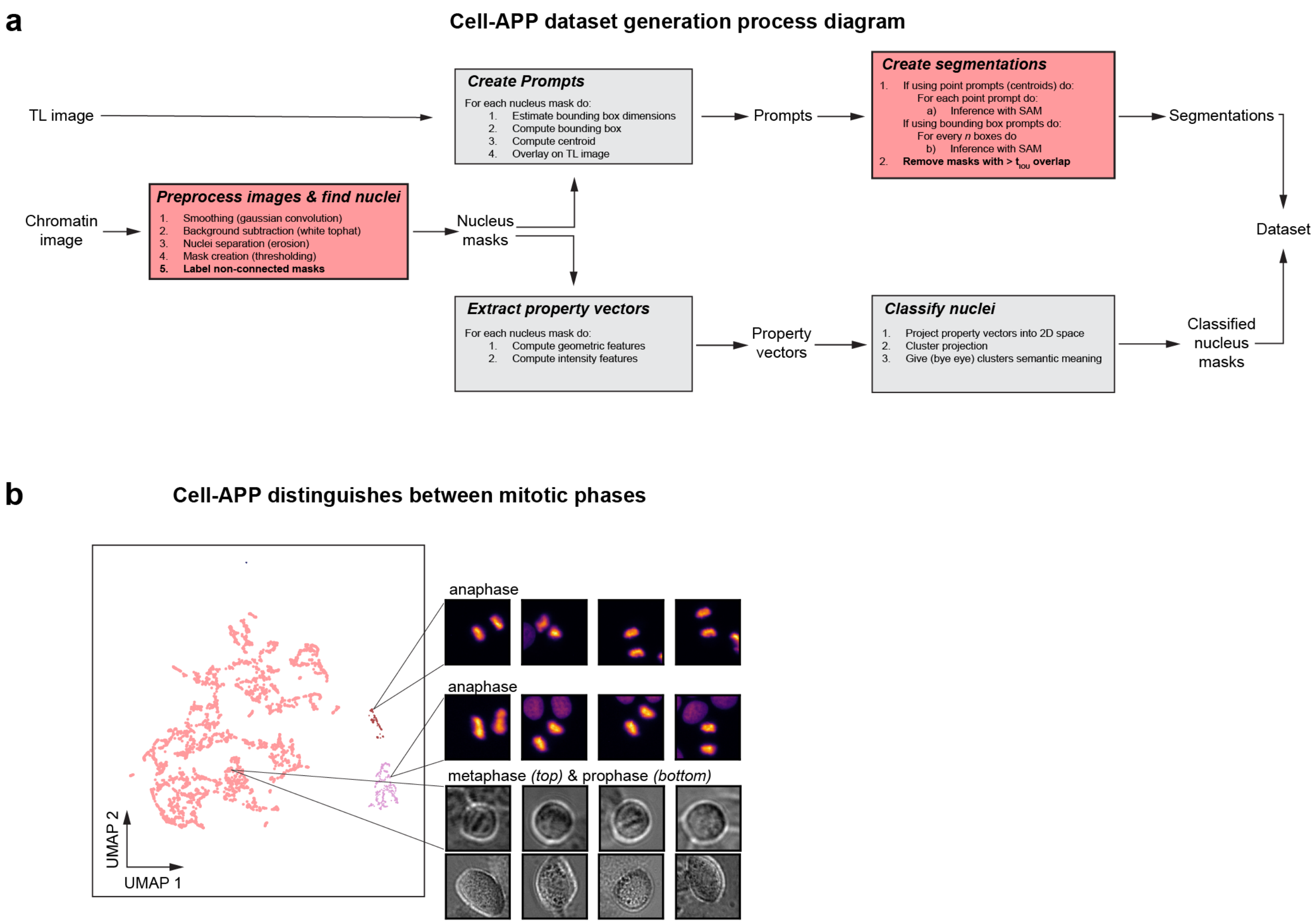


**Figure S1**. **Dataset generation process diagram. a,** Diagram mirrors **Fig. 1**. Unboxed texts represent objects; boxed texts represent functions. Boxes shaded red indicate that the function may "lose" annotations. *Preprocess images & find nuclei* may not find all nuclei if nuclei are touching. Cells not expressing the chromatin fluorescent tag will be "lost" too. In *Create segmentations,* after SAM generates all segmentations for a given image, any segmentations that overlap significantly are discarded. Significant overlap is defined by an IOU greater than *t,* where *t* is a user-defined tolerance. Lost annotations will reduce dataset quality and ultimately influence model performance. **b,** Anaphase nuclei can be found by re-projecting and clustering the mitotic cluster from **Figure 1b**. We note, however, that these anaphase clusters are contaminated with minimal dead, prophase, and metaphase cells.

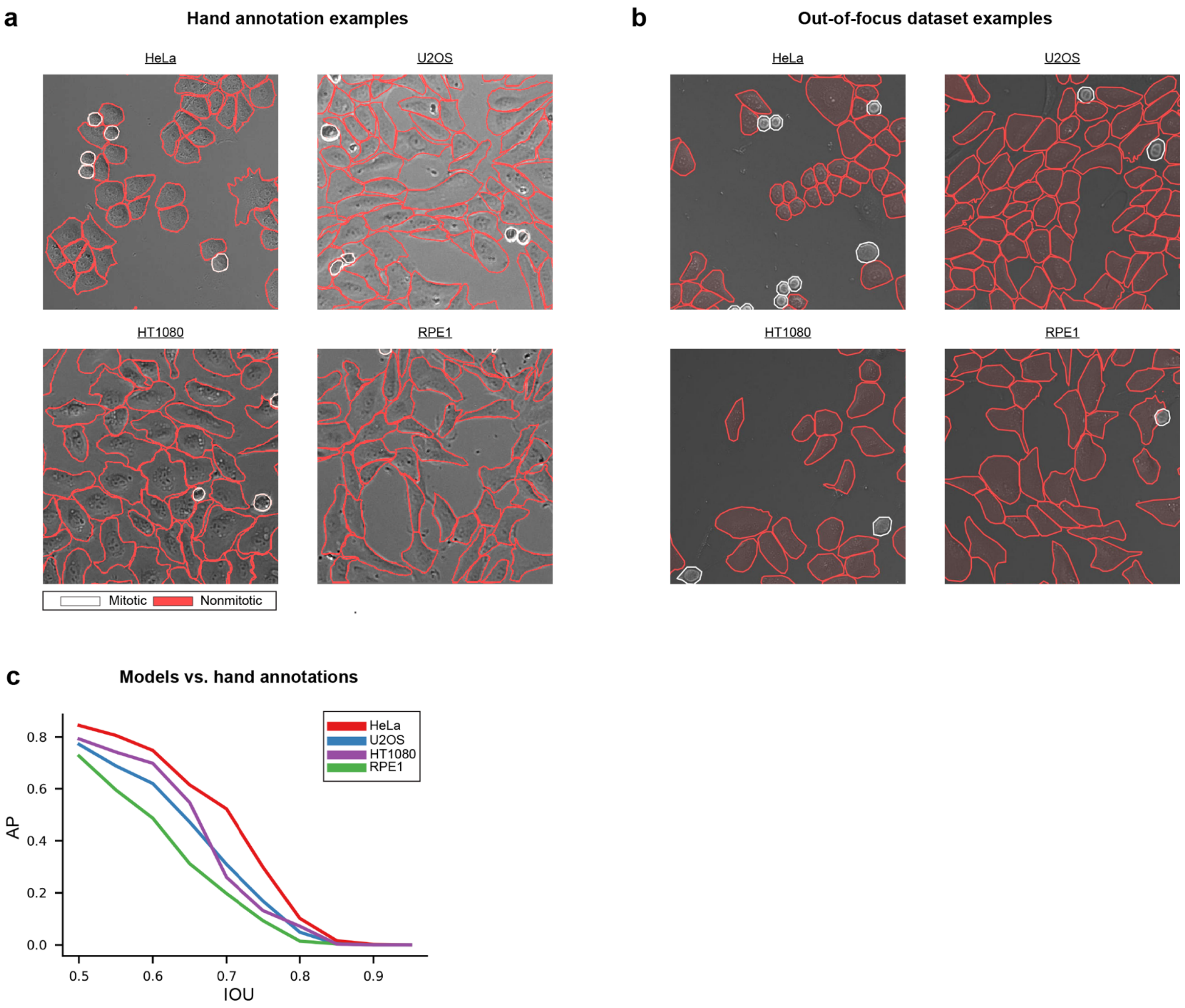


**Figure S2. Additional annotation examples. a,** Examples of the hand annotations used in **Fig. 2b. b,** Examples of out-of-focus images and annotations. Images from "three" qualitatively distinct focal planes—above focus, focus, and below focus—were used to train each model. "Above" and "below" focus images are considered "out-of-focus". **c,** Evaluating cell line-specific models on hand annotations (of the same cell line) reveals that differences in model performance are less drastic than **Figure 3c** suggests. This confirms that dataset quality influences model performance metrics.

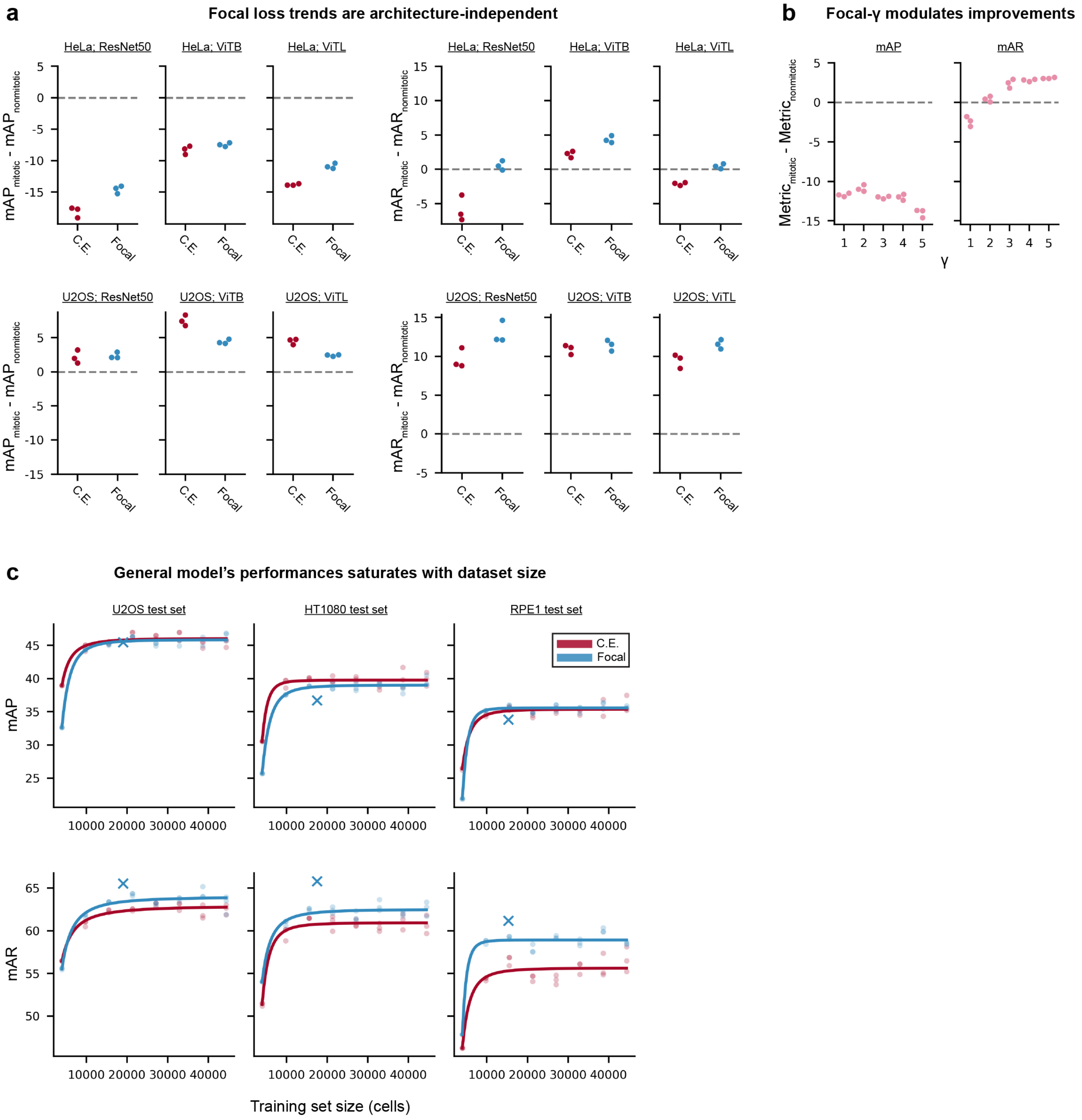


**Figure S3. Additional properties of focal loss and the general models. a,** The ways in which focal loss modulates models' performance on mitotic cells, with respect to nonmitotic cells, are independent of model architecture. **b,** Focal loss improves average precision on mitotic cells, only for $\gamma < 2$; however, it improves mAR for all $\gamma$. **c,** Evaluation of the general models on the U2OS, HT1080, and RPE-1 testing datasets shows that performance saturates after dataset size surpasses ~10k cells.

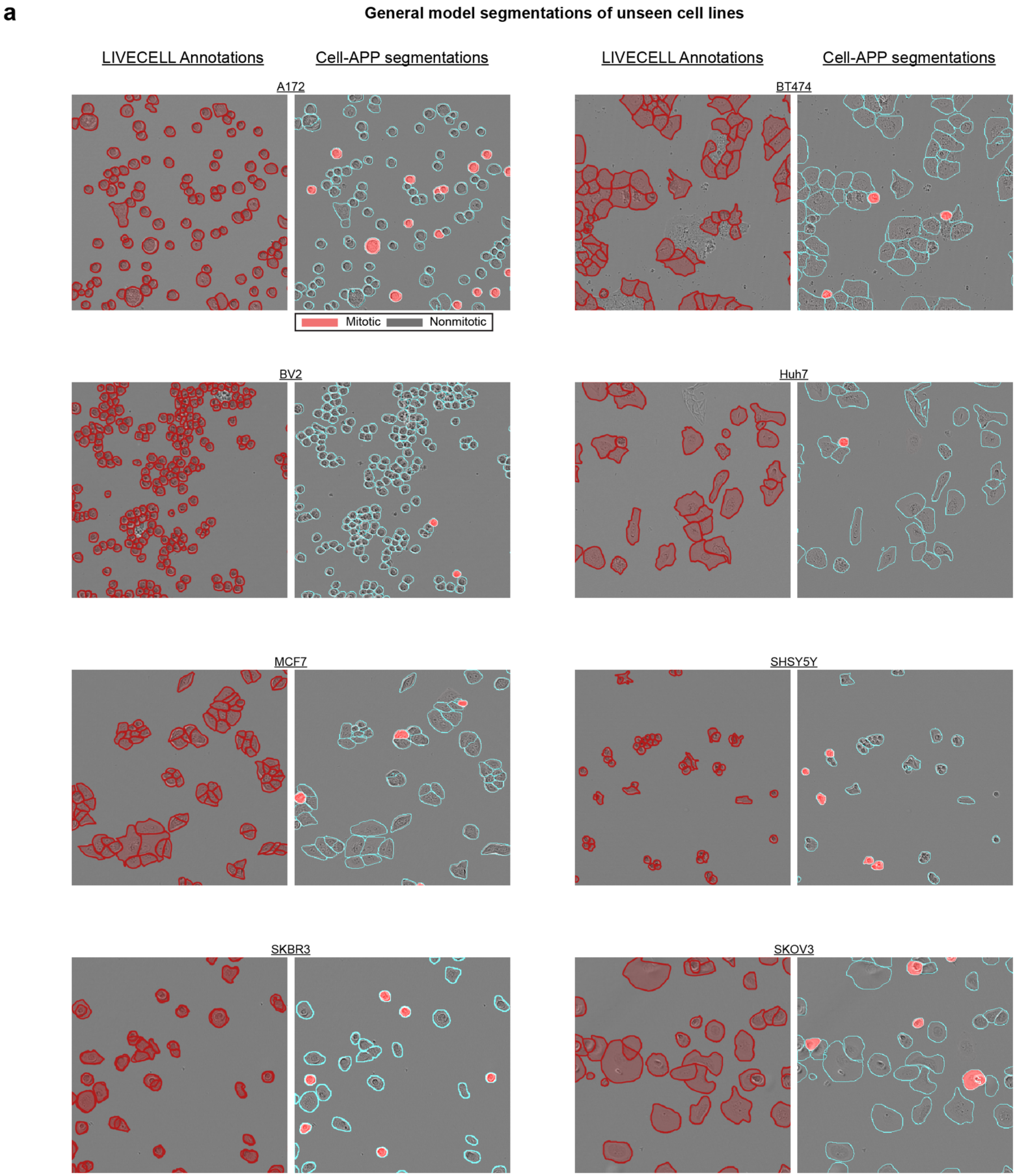


**Figure S4. The Cell-APP general model can segment unseen (OOD) cells. a,** *Left:* Hand-annotations from the LIVECELL dataset. *Right:* Segmentations produced by a Cell-APP model trained on the general dataset containing ~45k cells equally distributed across HeLa, U2OS, HT1080, and RPE1 cell types.

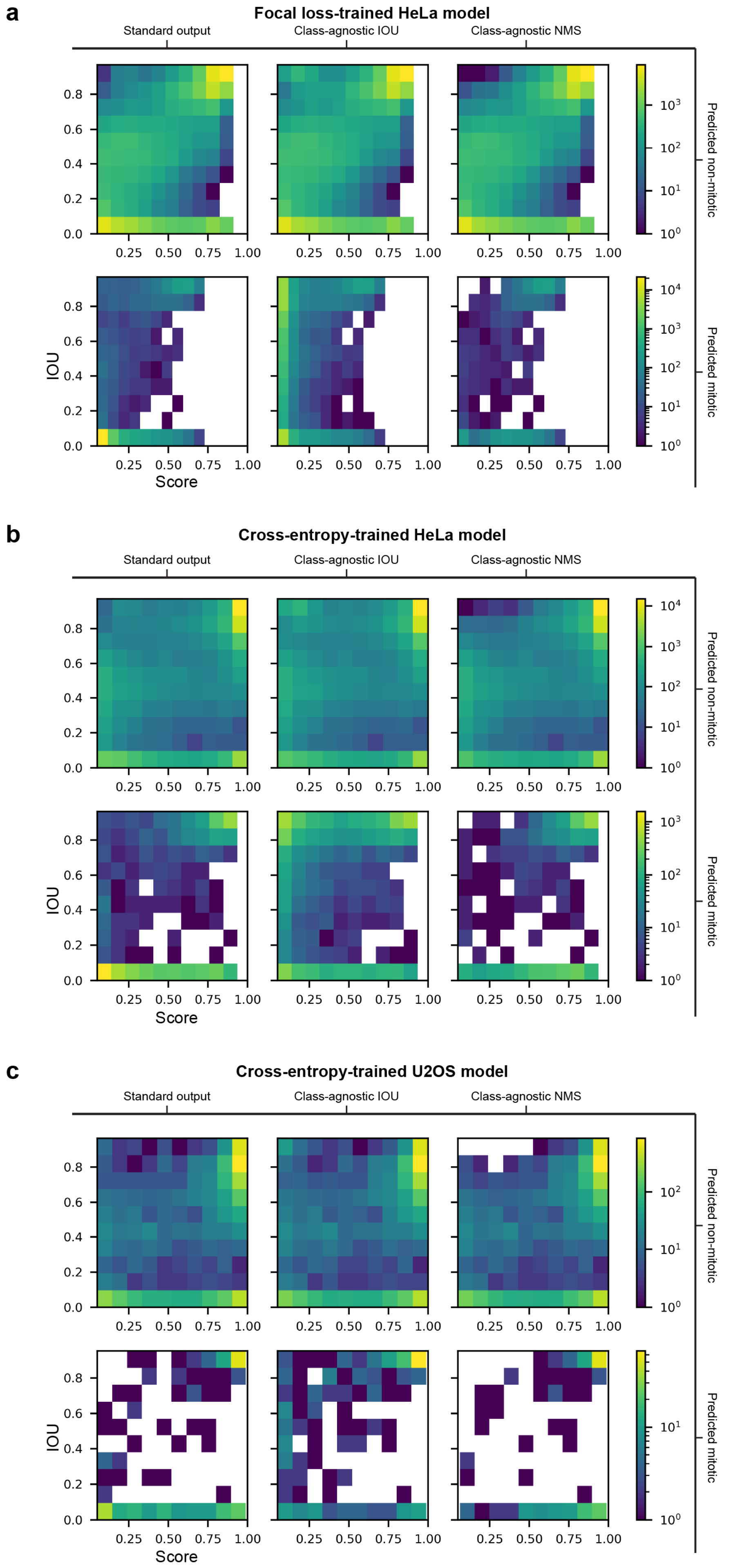


**Figure S5. Common model errors.** All plots depict joint distributions for the score and IOU of a model's predictions on either the HeLa (**a,b**) or U2OS (**c**) test set. Score describes the model's confidence in a given prediction; IOU describes a given prediction's relative overlap with the nearest annotation in the test set. In the left-most histograms, misclassifications have an IOU = 0 (i.e., if the model predicts a mitotic mask and the annotation is non-mitotic, IOU will always equal zero). This is standard COCO evaluation behavior. In the middle histograms, misclassifications can have an IOU > 0. In the right-most column, misclassifications have an IOU = 0, and if any masks are overlapping, the one with the lowest score is removed using non-max suppression. **a,** Distributions of the focal loss-trained HeLa model's predictions on the HeLa test set. By allowing misclassifications to have IOU > 0 *(middle),* we reveal that the focal loss-trained HeLa model misclassifies nonmitotic cells as mitotic more frequently than it misclassifies mitotic cells and nonmitotic. By applying non-max suppression, we show that most of the misclassifications (cells that gained IOU in the middle column; bottom left bin) are double predictions (i.e., the second predicted mask for a given cell). Applying non-max suppression also reveals that the focal loss-trained HeLa model makes correctly classified double predictions (compare top-left corner densities in the left and right column plots). **b,** Distributions of the cross-entropy loss-trained HeLa model's predictions on the HeLa test set, showing these trends are loss-function invariant. **c,** Distributions of the cross-entropy loss-trained U2OS model's predictions on the U2OS test set, showing these trends are cell line invariant.

| Geometry-based | Intensity-based |
|---|---|
| area | maximum intensity |
| solidity | minimum intensity |
| crofton perimeter | standard deviation of intensity |
| convex hull area | mean intensity |
| eccentricity | |
| major axis length | |
| minor axis length | |
| perimeter | |
| centroid | |
| euler number | |
| extent | |
| filled area | |

**Table 1.** Geometric and intensity-based features used in the nuclei classification pipeline.

| | $N_{mitotic}$ | $N_{nonmitotic}$ | Median Area$_{mitotic}$ | Median Area$_{nonmitotic}$ | Median Annotations per Image | Median Confluency |
|---|---|---|---|---|---|---|
| HeLa | 4311 | 58241 | 222.4 | 356.1 | 139 | 25.00% |
| U2OS | 755 | 18300 | 245.5 | 518.1 | 244 | 58.60% |
| HT1080 | 638 | 16873 | 304.9 | 565 | 78 | 28.00% |
| RPE1 | 495 | 14785 | 265.3 | 711.8 | 86 | 28.70% |

**Table 2.** Summary statistics on the cell line-specific datasets.

| Dataset | $N_{mitotic}$ | $N_{nonmitotic}$ | $N_{HeLa}$ | $N_{U2OS}$ | $N_{HT1080}$ | $N_{RPE1}$ | Median Annotations per Image | Median Confluency |
|---|---|---|---|---|---|---|---|---|
| 1 | 166 | 3789 | 996 | 965 | 994 | 1000 | 47 | 25.50% |
| 2 | 501 | 9145 | 2438 | 2325 | 2441 | 2442 | 75 | 36.00% |
| 3 | 769 | 14670 | 3882 | 3803 | 3870 | 3884 | 86 | 42.00% |
| 4 | 1115 | 20179 | 5327 | 5316 | 5325 | 5326 | 82 | 39.00% |
| 5 | 1418 | 25502 | 6767 | 6623 | 6762 | 6768 | 88 | 44.10% |
| 6 | 1646 | 31115 | 8210 | 8144 | 8206 | 8201 | 83 | 41.20% |
| 7 | 1986 | 36610 | 9650 | 9643 | 9650 | 9653 | 86 | 42.40% |
| 8 | 2281 | 42057 | 11095 | 11058 | 11088 | 11097 | 87 | 41.50% |

**Table 3.** Summary statistics on the multi-cell line datasets.

| | *mAP* | *mAR* |
|---|---|---|
| *HeLa-CE* | 57.1 ± 0.337 | 72.7 ± 0.132 |
| *HeLa-focal* | **59.0 ± 0.326** | **74.8 ± 0.342** |
| *General (8)-CE* | 57.6 ± 0.309 | 72.2 ± 0.675 |
| *General (8)-focal* | 57.2 ± 0.203 | 73.1 ± 0.225 |

| | *mAP* | *mAR* |
|---|---|---|
| *HT1080-CE* | 36.4 ± 0.167 | 56.8 ± 0.482 |
| *HT1080-focal* | **36.8 ± 0.177** | **61.8 ± 0.729** |
| *General (8)-CE* | 36 ± 1.04 | 56.6 ± 1.2 |
| *General (8)-focal* | 35.7 ± 0.164 | 58.5 ± 0.134 |

| | *mAP* | *mAR* |
|---|---|---|
| *U2OS-CE* | 45.2 ± 0.661 | 61.2 ± 0.244 |
| *U2OS-focal* | 44.0 ± 0.224 | 62.9 ± 0.338 |
| *General (8)-CE* | 45.4 ± 0.511 | 62.5 ± 0.482 |
| *General (8)-focal* | **46.4 ± 0.507** | **63.1 ± 0.888** |

| | *mAP* | *mAR* |
|---|---|---|
| *RPE1-CE* | 31.1 ± 0.214 | 54.0 ± 1.04 |
| *RPE1-focal* | 33.5 ± 0.862 | 57.7 ± 0.877 |
| *General (8)-CE* | 38.9 ± 0.840 | 60.7 ± 0.874 |
| *General (8)-focal* | **39.6 ± 0.667** | **62.6 ± 0.680** |

**Table 4.** Summary of model performance. Each quadrant denotes a different test set (top left: HeLa, top right: HT1080, bottom left: U2OS, bottom right: RPE1). The left most column of each quadrant denotes the model. Each metric is the mean ± standard deviation from three best performing sets of weights (where the different weights are from different training epochs).
